# Acute and chronic pesticide exposure trigger fundamentally different molecular responses in bumble bee brains

**DOI:** 10.1101/2024.10.09.617417

**Authors:** Alicja Witwicka, Federico López-Osorio, Andres Arce, Richard J Gill, Yannick Wurm

## Abstract

Beneficial wild insects including pollinators encounter various pesticide exposure conditions, from brief high concentrations to continuous low-level exposure. To effectively assess the environmental risks of pesticides, it is critical to understand how different exposure patterns influence their effects. Unfortunately, this knowledge remains limited. To clarify whether different exposure schemes disrupt the physiology of pollinators in similar manners, we exposed bumble bees to acute and chronic treatments of three different pesticides: Acetamiprid, clothianidin, and sulfoxaflor. Gene expression profiling enabled us to compare the effects of these treatments on the brain in a high-resolution manner. There were two main surprises: First, acute and chronic exposure schemes affected largely non-overlapping sets of genes. Second, different pesticides under the same exposure scheme showed more comparable effects than the same pesticide under different exposure schemes. Acute exposure caused up-regulation of stress response mechanisms causing distinct regulatory changes, rather than amplifying the effects of prolonged low-dose exposure that affected predominantly immunity and energy metabolism. These findings show that the mode of exposure critically determines the effects of pesticides. Our results signal the need for safety testing practices to better consider mode-of-exposure dependent effects and suggest that transcriptomics can support such improvements.

## Introduction

Safeguarding insect pollinators is critical for ecosystem stability, food production, and human welfare (Potts et al., 2016). Agricultural pesticides significantly contribute to pollinator declines, including through indirect sublethal effects (Siviter et al., 2021). These declines have prompted calls for a better understanding of how different exposure schemes affect physiology and behavior (Gill & Raine, 2014; Kenna et al., 2019). Indeed, exposure in natural environments varies from low to high concentrations over variable periods, potentially challenging the accuracy of risk assessments (EFSA, 2023) and our ability to forecast population responses. However, the extent to which different exposure schemes have distinctive molecular and physiological effects is unknown.

Focusing assessment of pesticide toxicity on few selected phenotypes can result in contradictory measures of their negative effects (Tsvetkov & Zayed, 2021). In contrast, advances in transcriptomics enable us to measure genome-wide effects of pesticide exposure (Grozinger & Zayed, 2020; López-Osorio & Wurm, 2020). Indeed, simultaneously quantifying expression levels of thousands of genes provides high-dimensional data that enables assessment of many aspects of insect physiology. Applying the higher-resolution approaches could enable new insights on the impacts and risks of pesticide exposure.

Here, we aimed to test whether different pesticide exposure schemes disrupt physiology of a beneficial pollinator species in similar manners. For this, we exposed microcolonies of *Bombus terrestris* bumble bees to “acute”, *i.e.*, short-term high-concentration and “chronic”, *i.e.*, long-term low-concentration pesticide treatments, and measured activity levels of all genes. We used microcolonies to increase power and control for genetic backgrounds. To investigate a range of outcomes, we performed experiments with three widely used pesticides: The neonicotinoid pesticides clothianidin and acetamiprid and the sulfoximine pesticide sulfoxaflor (Figure 1A-C). We measured brain gene expression because each of these pesticides target nicotinic acetylcholine receptors (nAChRs) which are common in the brain (Witwicka et al., 2023), and because the brain governs behavioral responses. We hypothesized that exposure to different pesticides would lead to distinct gene expression responses, and that acute exposure to a pesticide would primarily affect the same genes as chronic exposure, but with a greater intensity.

**Figure 1.**
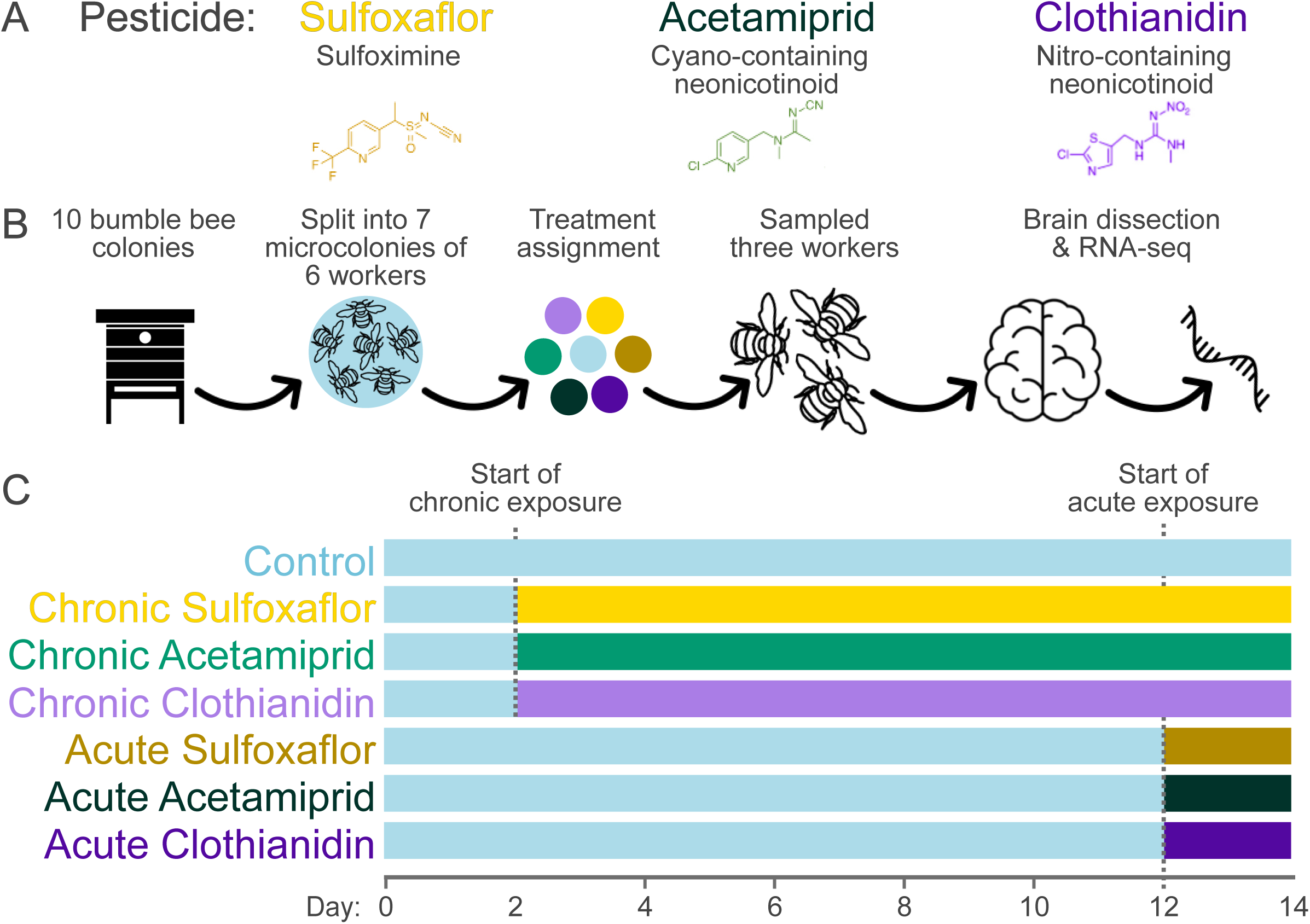
Experimental design. (A) Microcolonies were exposed to a control treatment or to one of three pesticides: Sulfoxaflor, acetamiprid, and clothianidin. (B) Seven microcolonies of six callow workers were obtained from each of the ten bumble bee source colonies and assigned to one of the seven treatments; RNA extractions focused on pooled brains of three workers per microcolony. (C) After a two-day adjustment period, we began chronic exposure; acute exposure began after 12 days to control for age of workers and day of sampling.

## Results

### Acute exposure causes stronger and broader changes than chronic exposure

We exposed bumble bee microcolonies to acute treatments (21.5 ppb, 48 hours) and chronic treatments (4.4 ppb, 12 days) of acetamiprid, clothianidin, and sulfoxaflor, and the control, and sequenced RNA from pools of brains of three workers per microcolony (Figure 1A-C). Acute exposure was always more disruptive than chronic exposure, resulting in greater numbers of differentially expressed genes. For example, acute exposure to clothianidin resulted in 3.5 times more differentially expressed genes than chronic exposure (Figure 1A and 2A). Similarly, changes in expression amplitude among the 20 genes with the most pronounced changes after acute exposure were 2.7 times greater than after chronic exposure (t-test, p < 10^-6^). In line with these patterns, principal component analyses showed that the expression profiles of chronically exposed microcolonies were more similar to control colonies than to acutely exposed colonies (Figure 2B).

**Figure 2.**
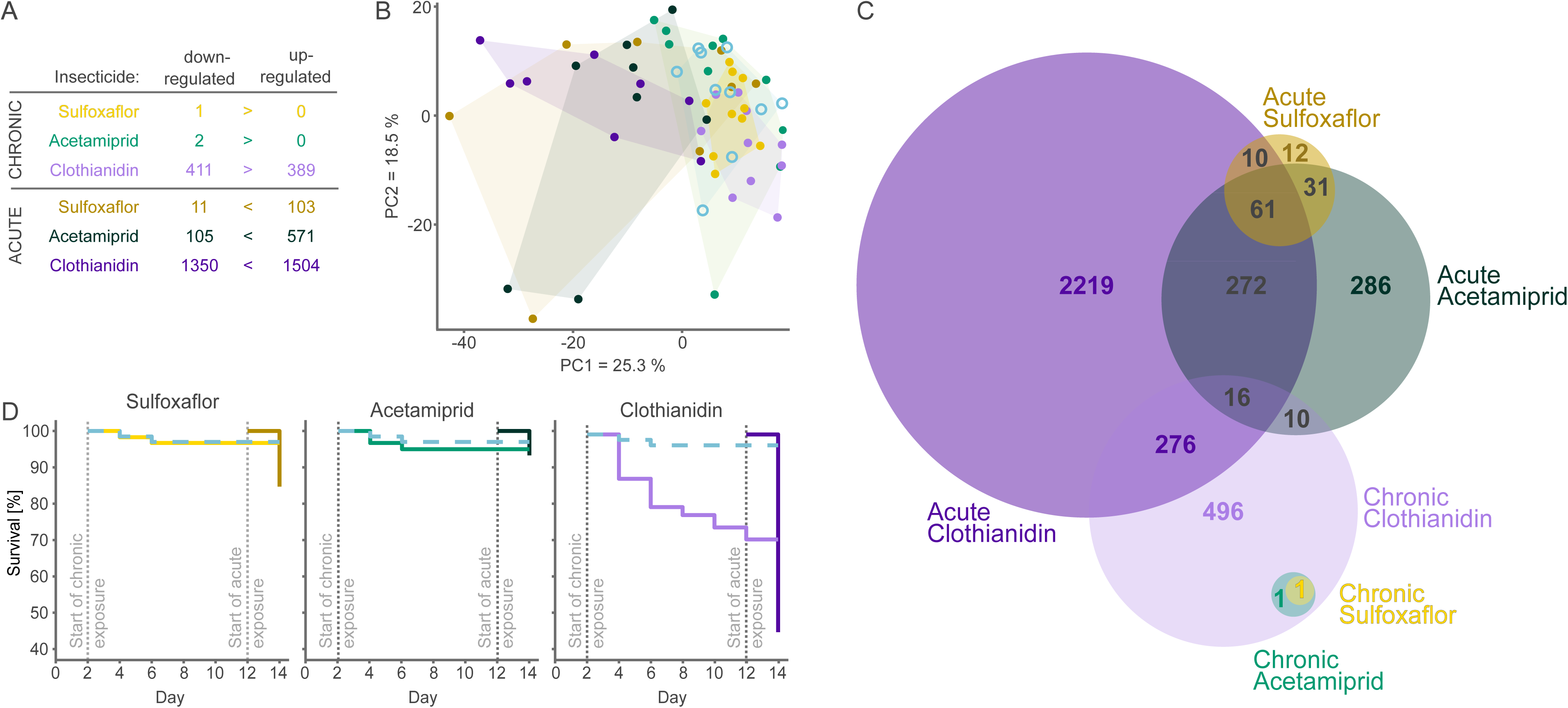
Gene expression changes and survival rates under chronic and acute exposure schemes. (A) Numbers of genes differentially expressed in each treatment compared to the control. (B) Principal Component Analysis of all microcolonies. The first principal component separates chronic and acute treatments; for the first two principal components, chronic treatments overlap with the control (blue). (C) Euler diagram showing relative numbers of differentially expressed genes and overlaps between treatments (DESeq2 Wald test cut-off of FDR 0.05). (D) Survival rates; Dashed blue lines indicate the control.

The effects of chronic exposure were also more consistent across the microcolonies than the effects of acute exposure (20% lower standard deviation; GLMM p < 10^-16^). The difference in variation may be because the organism’s physiology has more time to adjust over the 12 days of chronic exposure. Alternatively, genetic variation between colonies may affect the thresholds that trigger a response, as well as the rate at which the pesticide is taken up by the hemolymph and reaches the brain (Figure S1 and S2).

### Different exposure schemes drive different types of changes

There were major differences in the magnitude of effects between pesticides: Acute exposure to clothianidin and acetamiprid respectively changed the expression of 25 and 6 times more genes than acute exposure to sulfoxaflor. Chronic exposure to clothianidin similarly caused substantially more changes than acetamiprid or sulfoxaflor (Figure 2A). Nevertheless, we observed discrepancies in the molecular responses between acute and chronic treatments. Expression levels of a set of 61 genes changed consistently in response to all acute treatments (40 times more than expected by chance, SE = 0.015; hypergeometric test, p < 10^-5^, Figure 2C). Moreover, 89% of genes differentially expressed under acute exposure to sulfoxaflor were differentially expressed in at least one other acute treatment. However, none of these overlapping genes were differentially expressed in chronic exposure treatments.

Although chronic exposure affected considerably fewer genes than acute exposure, the effects of different pesticides also overlapped (Figure 2A). In particular, expression of the antimicrobial peptide *defensin* was on average 32-fold lower in chronic treatments, while expression of another antimicrobial peptide, *abaecin*, was 11-fold lower after exposure to clothianidin and acetamiprid (Figure 3B). The other genes most significantly affected by chronic clothianidin exposure included the pathogen response genes *kynurenine/alpha-aminoadipate aminotransferase* and *antichymotrypsin-2*, which were respectively down-regulated 10-fold and 3-fold relative to the controls (Table S1). Changes in expression levels of immune genes may explain the decreased immunocompetence of bees exposed to neonicotinoids Intriguingly, expression of these immune genes was unaffected by acute exposure.

**Figure 3.**
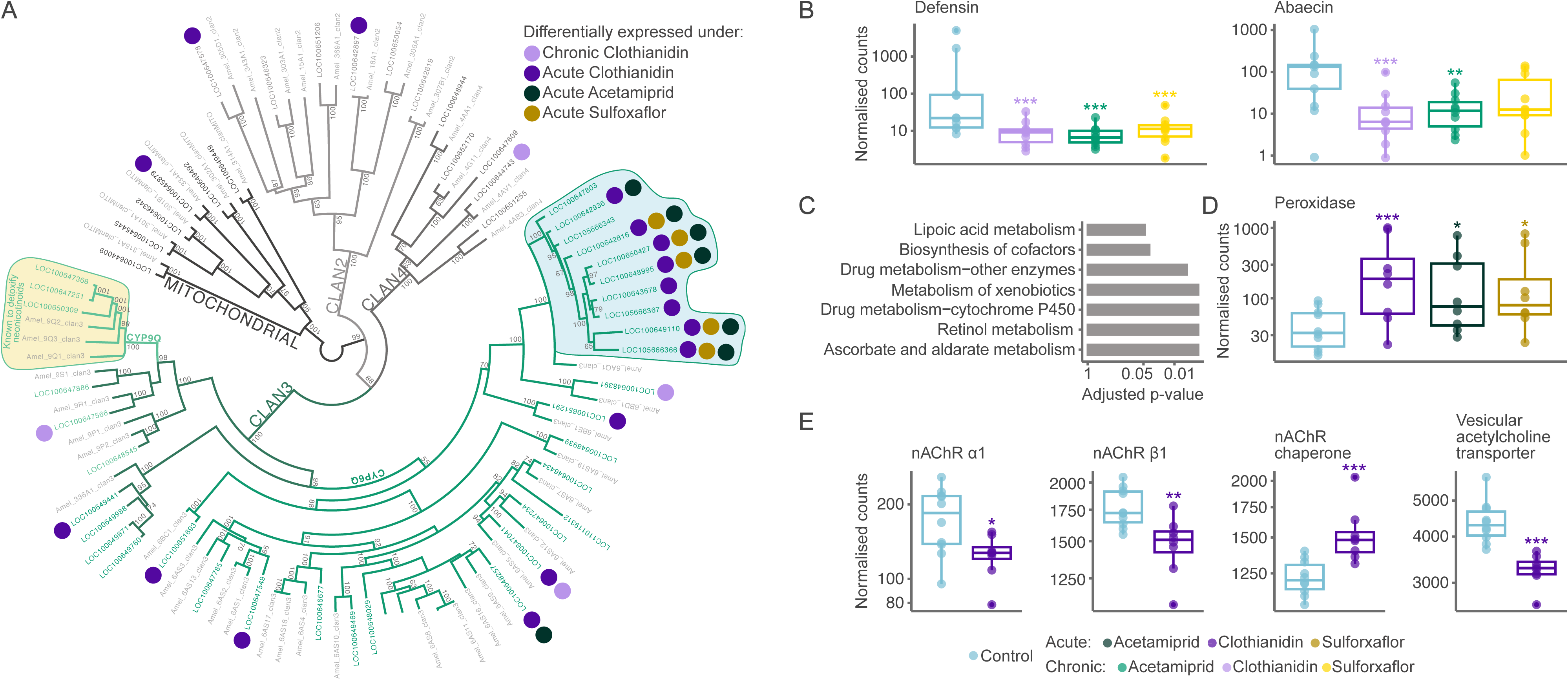
Selected genes affected by chronic and acute exposure schemes. (A) Maximum-likelihood tree of all cytochromes P450 in bumble bee (51 genes) and honey bee (49 genes). Filled circles highlight differentially expressed genes. Yellow area highlights CYP9Q genes known to detoxify neonicotinoids (Haas et al., 2022); none of these were up-regulated. Blue area highlights genes consistently differentially expressed under acute exposure. Numbers indicate bootstrap values of nodes. (B) Antimicrobial peptide genes were downregulated under chronic treatments. (C) KEGG pathways enriched in genes differentially expressed under all acute treatments. (D) *Peroxidase* was up-regulated in all acute treatments. (E) After acute clothianidin exposure, two nAChR-subunits whose *Drosophila* orthologs have particularly high affinity to clothianidin are downregulated, as is the vesicular acetylcholine transporter, while a chaperone which assists nAChR folding is up-regulated. Boxplot significance codes: * < 0.05; ** < 0.01; *** < 0.001.

### Acute exposure to every pesticide caused up-regulation of detoxification and stress-response genes

Among the 61 genes differentially expressed by all acute treatments, 91% were up-regulated rather than downregulated (*X^2^* = 23.8, p < 10^-6^). These included well-known detoxification genes (*e.g.,* P450 cytochromes, ATP-binding cassette transporters, and UDP-glycosyltransferases). Surprisingly, none of the cytochrome P450 genes from the CYP9Q subfamily were upregulated. Genes in this subfamily are conserved across bee species, and can determine neonicotinoid sensitivity (Haas et al., 2022; Manjon et al., 2018; Troczka et al., 2019). The three bumble bee CYP9Q genes most able to metabolize neonicotinoids have high baseline expression in the brain (representing 33% of the total expression of 51 cytochrome P450s; *X^2^ =* 21.5, p < 10^-6^). Perhaps there are constraints on the organism’s ability to further increase expression of these genes.

In contrast, expression of a subset of bumble bee lineage-specific cytochrome P450 genes in the CYP6Q subfamily increased by an average of 248-fold in acute treatments (Figure 3A). The dramatic up-regulation of these genes may help metabolize the pesticides or their metabolites, or it may contribute to a generic stress response. Indeed, other stress-response components included upregulation of ascorbic and lipoic acid metabolism and *peroxidase* (Figure 3C and D) likely indicating responses to oxidative stress which results from calcium ion influx after nAChR overstimulation (Hermann et al., 2015; Wang et al., 2018). Furthermore, acute exposure to acetamiprid and clothianidin caused up-regulation of peptide and fatty acid metabolism. This up-regulation may help compensate for the detrimental impacts of cellular stress, as indicated by analogous changes in mice (Yan et al., 2020).

### Acute and chronic clothianidin exposure schemes trigger mortality through different mechanisms

Although we primarily compared the effects of insecticides on bumble bee brains, we also measured survival, enabling us to examine the links between gene expression and mortality. Acute and chronic exposure to the most toxic compound, clothianidin, led to the death of 54% and 29% of bees, respectively (Cox proportional hazard models; both p < 10^-6^; Figure 2D). The cumulative doses of clothianidin consumed under both exposure schemes were comparable. This indicates that the effects of clothianidin do not cause stronger effects over time, and that bees are better able to tolerate long-term lower-level exposure than acute exposure.

Both acute and chronic clothianidin treatments affected a set of 292 overlapping genes, more than expected by chance (hypergeometric test, p < 10^-6^). However, most of the affected genes (90% and 64%, respectively) were treatment specific. Intriguingly, acute exposure up-regulated apoptosis-inducing genes, suggesting immediate lethal effects, while such links were absent after chronic exposure. Acute exposure to clothianidin also caused changes in the expression of genes important for nerve function, including up-regulation of synaptic vesicle proteins involved in neurotransmitter transport and release, and down-regulation of neuroligins which are crucial for synapse stability.

Acute clothianidin exposure also down-regulated nAChR subunits α1 and β1, and the vesicular acetylcholine transporter, and up-regulated the acetylcholine receptor chaperone which supports assembly of nAChRs (Figure 3E). Remarkably, α1 and β1 nAChR subunit orthologs in the fruit fly have particularly high affinity to clothianidin (Lu et al., 2022). These changes in gene expression suggest that some nAChRs in the brain may be reassembled after acute exposure, potentially to prevent further toxic effects. Different changes stood out after chronic exposure to clothianidin, including down-regulation of genes associated with the respiratory transport chain and ATP synthase complex (Fisher’s exact test, all p-values < 0.005). These findings parallel changes in locomotor behaviors whereby acute exposure increases locomotor activity, potentially because of synaptic overstimulation, whereas chronic exposure decreases locomotor activity, likely due to changes in energy metabolism (Kenna et al., 2019; Suchail et al., 2001).

## Discussion

Contrary to our initial hypothesis, the pesticide exposure scheme was a stronger determinant of bumble bee brain gene expression profiles than the type of pesticide. Indeed, we found more overlap among affected genes after acute exposure to different pesticides than between acute and chronic exposure to individual pesticides. This indicates that despite differences between pesticides in baseline toxicity levels and in chemical structures, the effects of exposure are strongly determined by exposure duration and intensity. Moreover, these pesticides do not show time-reinforced toxicity; the strength of effects does not increase over time. Instead, the effects differ altogether. Acute treatments cause severe and immediate stress responses, while chronic exposure gradually impairs immunity and energy metabolism. Thus, different exposure schemes pose distinct challenges for pollinators.

In agricultural settings, beneficial pollinators may survive pesticide exposure but suffer from sub-lethal disruptions that jeopardize their long-term abilities to reproduce and to pollinate crops and wild plants. Estimating the risks of such diverse detrimental impacts is essential for ensuring the environmental safety of pesticides. Traditionally, the acute LD_50_ dose which kills 50% of individuals over 24 or 48 hours has often been used as a proxy from which to draw conclusions about overall toxicity in other exposure scenarios (OECD Test no. 2013, 1998; OECD Test no. 214, 1998). Our findings make it difficult to justify such extrapolation because the manner and degree to which acute and chronic exposure schemes affect an organism at the molecular level differ substantially.

Recent revisions to the European Union’s testing guidelines include chronic 10-day exposure to pesticides and measurements of changes in coordination, hyperactivity, and shivering (OECD Test no. 245, 2017). Such improvements that consider sub-lethal effects should be applauded. However, the revised guidelines are likely unable to detect the diverse types of changes recognized by that gene expression profiling.

In the early days of examining gene expression to human diseases, only little information existed to help link changes in individual genes to tangible health outcomes (Hong et al., 2020). Similarly, we cannot yet directly link changes in gene expression to the ability of pollinators to survive, reproduce or pollinate. However, based on our understanding of cellular processes, changes in the expression levels of many genes undoubtedly indicate substantial disruption to basic biological functions.

Our findings show how measuring expression levels of thousands of genes can reveal previously unknown impacts of pesticides on beneficial species. Our work also highlights the complexities of such effects. We focused on two neonicotinoids and a sulfoximine which all target nicotinic acetylcholine receptors (nAChRs). The use and diversity of nAChR-targeting pesticides continue to increase (Goulson, 2013; Siviter & Muth, 2020), and we look forward to work clarifying how the patterns we observed here extend to pesticides with different modes of action, to other stressors, and in other beneficial insect species. Broader use of high-resolution approaches could significantly expand our understanding of how environmental stressors impact insect health. Just as gene expression profiling revolutionized the diagnosis and understanding of human disease, we anticipate it will play a pivotal role in the development and regulatory evaluation of novel pesticides.

## Acknowledgments

We were funded by the Natural Environment Research Council (NE/L00626X/1 and NE/S007229/1), the European Commission (H2020-MSCA-IF-2018-840185), and the Biotechnology and Biological Sciences Research Council (BB/T015683/1). This research used Queen Mary’s Apocrita HPC facility, supported by QMUL Research-IT (http://doi.org/10.5281/zenodo.438045).

## Data and code availability

Sequence data underlying this work are in the National Center for Biotechnology Information Short Read Archive, accession PRJNA1076820 (available upon publication). Analysis scripts will be available on GitHub at https://github.com/wurmlab/2022-09-Bter-chronic-acute-exposure-alicja.

During review, analysis scripts can be found at https://www.dropbox.com/scl/fo/h547ulwkbkc80zs4geyfm/h?rlkey=395la544cghli6ohqukdb6s91&dl=0

## Materials & Methods

### Pesticide solution preparation

Bumble bees were exposed to three pesticides that target nicotinic acetylcholine receptors (nAChRs): Clothianidin, acetamiprid or sulfoxaflor (Sigma Aldrich, UK). The use of clothianidin is restricted in the European Union (EFSA 2018), but it is still commonly used worldwide (Simon-Delso et al., 2015). Acetamiprid is the only neonicotinoid that can be applied in open fields without restrictions in the EU, with its approval renewed until 2033. Sulfoxaflor’s use has been increasing (Brown et al., 2016; Siviter et al., 2020) despite recent restrictions within the EU and reports of its high toxicity to bees when applied during flowering (EFSA, 2020). We used the same concentrations for each compound to allow for direct comparisons. We based the chronic concentration on the residues found in pollen and nectar (Botías et al., 2015, 2015; David et al., 2016; Pohorecka et al., 2012; Rundlöf et al., 2015; Siviter et al., 2018). We established the acute concentration based on the reports on clothianidin, the most toxic compound amongst these tested here to ensure survival of the bees and adequate replication levels for RNA sequencing. Reports suggest that 24h-LD_50_ for clothianidin varies greatly between colonies of *A. mellifera ligustica*, ranging from 9.69 ppb to 41.96 ppb (Laurino et al., 2013), we decided on a dose in the middle of that range. We established the chronic concentration at 5μg/L and the acute at five times higher, 25μg/L. We prepared pesticide stock solutions by dissolving pesticides in acetone to a concentration of 2.5mg/mL and 0.5 mg/mL and stored in darkness at −20L. Subsequently, we diluted the stock solutions using 30% sucrose solution to 5μg/L and 25μg/L feeding solutions. Given that the weight of a liter of 30% sucrose solution is 1130g, the final concentrations were 4.4 ppb and 22.1 ppb. To avoid pesticide degradation, we stored feeding solutions in darkness at 4L.

### Microcolony setup and sampling

We acquired ten source colonies of *Bombus terrestris audax* from a commercial breeder (Agralan Growers UK). We transferred the queen, existing brood and 20 workers to wooden boxes (30cm long/20cm wide/15cm deep) separated into two equal-sized chambers (foraging and nesting area). We provided each colony with *ad libitum* 30% sucrose solution and organic honeybee-collected pollen (General Food Merchants LTD). We marked all twenty workers using water-based markers (POSCA, UK) and screened the colonies daily for the emergence of new workers. All source colonies were two weeks old when we started setting up the microcolonies. Six callow workers that emerged within 24h were used to assemble microcolonies. Over 14 days, we obtained seven microcolonies from each source colony in a staggered manner, depending on the pace of worker production (Figure S3). Each microcolony was kept in a single-chamber wooden box (12cm long/12cm wide/10cm deep) and provided with organic pollen *ad libitum* and a single dose of nest substrate (5 parts organic pollen to 1 part 30% sucrose solution in a 2.5 cm Petri dish). Each microcolony received 10 mL of 30% control sucrose solution across two 5 mL syringes. We assigned the control treatment or one of the six pesticide treatments to all microcolonies in a randomized fashion. Our design ensured that treatments were assigned to microcolonies created at various stages of the source colony development. We replaced the control syringes with two syringes containing pesticide solution (a) 4.4 ppb after two days for chronic exposure or (b) 22.1 ppb after twelve days for short-term, acute exposure, so that all workers were of the same age at the time of sampling. We changed pollen and sugar solution every 48h between 11:00 am and 1:00 pm. All colonies remained in darkness at 24L, and we used red light during feeding and sampling. All sampling took place between 1:00 pm and 2:00 pm to minimize the effects of circadian rhythms on gene expression profiles. We placed individual workers in 2 mL screw-cap cryovial tubes, rapidly submerged them in liquid nitrogen, and stored the samples at −80L. We obtain ten replicates per treatment except for acute clothianidin were we obtained eight replicates because the mortality rates in two source colonies were continuously too high.

### RNA extraction, library preparation and high throughput sequencing

In the absence of a queen, one bumble bee worker can become dominant, which induces development of bigger ovaries. We examined abdomens to check for ovarian development and excluded dominant workers from further steps as significant ovarian development may change gene expression patterns. We dissected brains on dry ice and immediately placed them in a homogenization tube. We randomly selected three non-dominant workers per microcolony and pooled brains for further steps. We homogenized the dissected brains in TRIzol using a FastPrep96 (45 seconds at 1800 RPM). We isolated RNA using chloroform and purified it with Genaxxon RNA Mini Spin Kit, applying DNase I on-column digestion. We prepared cDNA libraries using the NEBNext® Ultra™ II Directional RNA Library Prep Kit for Illumina. We altered the standard protocol using one-third volumes of enzymes and buffers and 300 ng of total RNA input with 14 PCR cycles. Library size was quantified using TapeStation 2200 (Agilent, UK) and Qubit 2.0 fluorometer. Libraries were sequenced at QMUL Genome Centre on NextSeq 500 generating a mean of 36 million 40 bp paired-end reads per sample (from 22.8 million to 52.2 million reads).

### Quality assessment of raw reads

Initially, we assessed the quality of raw reads using FastQC v0.11.9 (Andrews, 2019). To evaluate alignment qualities, we respectively aligned RNA-seq samples to the *B. terrestris* genome and transcriptome (Ensembl Metazoa release 52) using STAR v2.7 (Dobin et al., 2013) and kallisto v0.46 (Bray et al., 2016). We processed STAR alignments using the RNA-seq module of Qualimap v2.2.1 (Okonechnikov et al., 2016) and summarized the results from FastQC and Qualimap using MultiQC v1.9 (Ewels et al., 2016). Four samples failed our quality-control checks due to poor alignment (<65%) and unusual GC content, which can result from PCR errors or contamination (Conesa et al., 2016). In total, we retained 64 samples, including 10 replicates for controls and all chronic treatments and 8 replicates for each acute treatment.

### Differential gene expression analysis

We summarized kallisto pseudo-aligned transcript abundances to the gene level with tximport v1.14 (Soneson et al., 2016) using transcript-to-gene tables retrieved with biomaRt v2.48.3. We excluded genes with low expression levels applying a cut-off of at least eight samples with a count of 10 transcripts or higher, retaining 9,735 out of 10,661 genes with mapped reads. We transformed counts using the variance stabilizing transformation (VST, *blind = FALSE*) and performed principal component analysis on all remaining genes to assess the relationships of the samples. To detect differentially expressed genes between exposure treatments and the control, we applied Wald tests on median-of-ratios normalized counts in DESeq2 v1.32.0 (Love et al., 2014). We included treatment and source colony as factors in the model design. We report genes as differentially expressed using a significance cut-off of 0.05 after false discovery rate adjustment (Benjamini & Hochberg, 1995) of the Wald test p-values (FDR).

### Gene Ontology enrichment analysis

To test which Gene Ontology terms were overrepresented in response to the treatments we performed gene ontology enrichment analysis using g:GOSt option of gprofiler2 v0.2.1 (Kolberg et al. 2020). We sorted differentially expressed genes (Wald test’s FDR < 0.05) for each treatment by adjusted p-values change values and set the custom background genes to all genes expressed in control brain samples. We report Gene Ontology terms as enriched applying corrected for multiple testing g:SCS threshold of 0.05 derived from Fisher’s exact test.

### Cytochrome P450 expression analysis and phylogeny

We identified 51 putative cytochrome P450 orthologs for *B. terrestris* searching the proteome for the Pfam PF00067.25 domain and extracted them using HMMER v3.1b2 applying the *--cut_ga* option, which removes domains with conditional E-values greater than the internally established threshold.

We selected the longest isoforms of the 51 *B. terrestris* and 49 *A. mellifera* P450 (Dermauw et al., 2020) cytochromes P450, and used the *mafft-linsi* option of MAFFT v7.480 (Katoh & Standley, 2013) to perform multiple sequence alignment of amino acid sequences. We trimmed the aligned sequences using Jalview v2.11 (Waterhouse et al., 2009) and TrimAl using *-automated1* option (Capella-Gutiérrez et al., 2009). We used IQ-TREE v2.0.3 (Nguyen et al., 2015) to perform maximum likelihood phylogenetic inference using the *-MFP* option (Kalyaanamoorthy et al., 2017).

### Variable number of replicates in treatment groups

To examine whether the statistical power of detection of the differentially expressed genes changes due of the uneven number of replicates across exposure groups (8 in acute treatments and 10 in chronic treatments), we ran the analyses again using eight replicates per treatment. We randomly selected 8 samples for the control and chronic treatment and repeated Wald test as implemented in DESeq2. We ran this analysis over 50 iterations making sure that the random exclusion of samples is unique for each iteration. We identified genes as differentially expressed applying a cut-off for false discovery rate (FDR) < 0.05. We built a distribution of the number of differentially expressed genes detected at each iteration to assess the probability of obtaining a higher number of differentially expressed genes for each of the acute treatments when the number of replicates is equal in all groups. All of the general trends we report here held under the alternate scenarios. Our analysis indicates that reducing the number of replicates in chronic treatments does not increase the power of detection of differentially expressed genes for acute treatments (permutation test, all p > 0.09). Therefore, we decided to retain all the suitable samples in downstream analysis to allow for a better estimation of the gene expression levels.

### Significance of overlaps of differentially expressed genes between treatments

To determine if the overlap between differentially expressed genes across various treatments was statistically significant, we used a simulation-based method, anchored in a hypergeometric testing framework. Specifically, we created 10,000 unique samples by randomly selecting genes from a combined pool of all expressed genes without replacing them once drawn. The number of genes drawn was determined by the actual number of differentially expressed genes under each treatment. For each comparison we assessed the extent of overlap between treatments and compared to the actual observed overlap in the real data. We calculated the p-value as the proportion of simulated overlaps that were at least as extreme as the observed overlap, adjusted for the observed case itself.

### Food intake measurements

Every other day, we measured food intake of each microcolony. We calculated the median pesticide dose [μg] per bee over exposure time for both acute and chronic exposures. Microcolonies exposed to acute clothianidin treatment, consumed a median of 0.017μg of clothianidin, while microcolonies exposed to clothianidin at chronic regime consumed 0.019μg. For acetamiprid, microcolonies consumed on average 0.036μg and 0.039μg of the compound under acute and chronic exposure respectively. Microcolonies exposed to sulfoxaflor consumed a median of 0.038μg and 0.040μg of the compound under acute and chronic exposure respectively.

We used a linear model to assess differences in the food intake between the control and chronic treatment groups. Specifically, we used the average daily food intake per bee within a microcolony as the response variable and included the queenright colony of origin and the interaction between day of exposure and treatment as explanatory variables. Our model returned a negative coefficient of −0.063 with a small standard error of 0.015 for an interaction term between the day of exposure and chronic clothianidin treatment, indicating that the effect of chronic clothianidin exposure on per day food intake decreased by 0.063 mL for each consecutive measurement compared to the control (p < 10-5). We observed a similar trend applying a linear model with the same variables to food intake data from microcolonies exposed to acute doses of clothianidin (coefficient = −0.51, SE = 0.11 p < 10-5). In this model we only used measurements from days 12 (last day before exposure) and 14 (sampling day post exposure). We did not detect significant differences in the food intake between the control microcolonies and microcolonies exposed to sulfoxaflor and acetamiprid at either acute or chronic treatments.

The bees exposed to clothianidin consumed less compound compared to sulfoxaflor and acetamiprid, however, even at a lower total dose this compound caused the most changes. Most importantly, the accumulative dose consumed under acute and under chronic exposures were comparable for all compounds. It is important to note that some of the changes detected in the differential gene expression analysis between control samples and bees exposed to chronic clothianidin may have been caused by lower food intake rather than the exposure alone. Unfortunately, we were unable to differentiate between these two factors. Nonetheless, the observed lower food intake was a direct result of the exposure to clothianidin.

### Quantification of differences between source colonies

To detect differentially expressed genes between the source colonies, we used the same DESeq2 model that was used for the detection of differentially expressed genes between treatments, which included treatment and source colonies as explanatory variables. We report genes as differentially expressed using a significance cut-off of FDR < 0.05 (Figure S1). After mapping reads obtained from all microcolonies to the reference genome using STAR, we mapped variants using freeBayes v1.3.1 (Garrison & Marth, 2012) and applied principal component analysis using SNPRelate v3.18 (Zheng et al., 2012) to see whether there are differences in the genetic makeup between the source colonies. All colonies were separated by the first two principal components showing that the source colonies varied in their genetic background (figure S2).

### Variable responses in bees exposed to acute pesticide treatments

We observed that the variance in gene expression between replicates in acute treatments is higher than in chronic and control treatments. To check if this pattern is statistically significant, we built a generalized mixed-effects model using lme4 R library. We used standard deviation of genes as a response variable and treatment (acute or chronic) and differential expression status (differentially expressed or unchanged) of each gene as fixed explanatory variables, we assigned gene as a random effect to account for a non-independence of the observations. To address issues related to heteroscedasticity and non-normality, we applied variance stabilizing transformation (VST) before calculating the standard deviation of gene expression. The use of VST helped to mitigate the presence of extreme values that were observed when standard deviation was calculated on un-normalized counts, which resulted in a skewed distribution of the residuals, and hindered our ability to fit a generalized linear model. Standard deviation of gene expression values after VST is more consistent across genes, as VST reduces the dependence of variance on the mean expression level. Therefore, if we still see differences in standard deviation between acute and chronic groups calculated using VST-normalized counts, the differences in standard deviation between genes in non-normalized counts should be greater. To assure the best model fit for continuous data with a right-skewed distribution, we applied gamma distribution.

The model’s output showed that the interaction term between treatment and the expression status variables has a significant effect on the response variable, with a coefficient of 0.19, t-value of −13.91, and p of < 10^-16^, suggesting standard deviation is higher for the acute group than for the chronic group when genes are differentially expressed. In the context of GLMM with a gamma distribution, the coefficient is on the log scale. Therefore, the standard deviation in the acute treatment group is approximately 20% higher on average compared to the chronic group.

### Survival analysis

We collected data on the survival of individual bees throughout the experiment. We fitted Cox proportional hazards regression models to conduct pairwise comparisons of survival between each treatment and the control utilizing *survfit* function from the survival v.3.5-7 R library. For chronic treatments, we compared the data from the entire experiment duration (14 days). For the acute treatments we only compared survival data from the last two days of the experiments when microcolonies assigned to acute treatments experienced exposure.

### Differences in the scale of treatment effects

To assess whether differentially expressed genes under acute exposure are more likely to exhibit substantial changes in expression levels compared to those under chronic exposure, we calculated the proportion of genes showing at least a 4-fold change in expression relative to the control. A Chi-square test was utilized to determine whether this proportion was significantly higher in the acute treatment group. The findings indicated that 3.61% of differentially expressed genes under acute clothianidin treatment experienced at least a 4-fold expression change, in contrast to only 0.75% of differentially expressed genes under chronic treatment (χ² = 16.7, p < 0.00001). Moreover, the most extreme expression change under acute exposure to clothianidin was a 1487-fold increase, which is substantially larger than the 31-fold decrease seen with chronic exposure. Thus, we selected 20 genes with the most extreme changes under acute and chronic clothianidin treatments and ran Welch two sample t-test to compare the log2Fold changes between the gene sets. On average, the top 20 affected genes had an increase of expression 2.65 times higher under acute clothianidin exposure compared to the chronic exposure (t = 6.2, df = 26.4, p < 10^-6^).

## Notes

### Competing Interest Statement

The authors have declared no competing interest.

