## Supplemental Information for "Acute and chronic pesticide exposure trigger fundamentally different molecular responses in bumble bee brains"

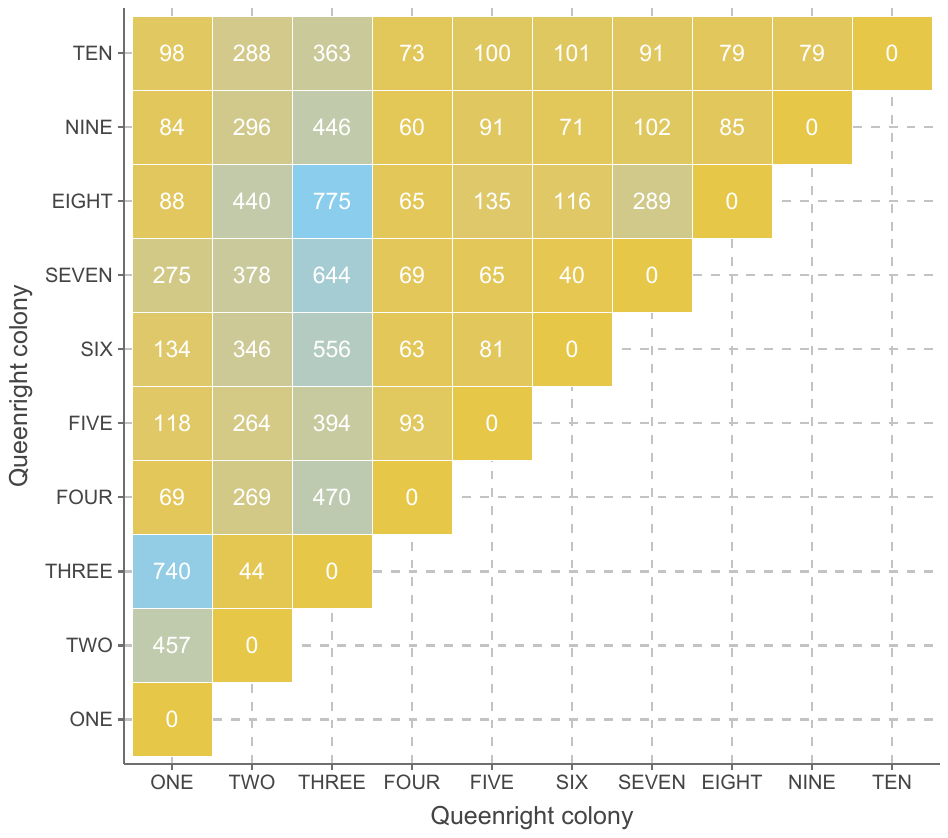

Figure S1. Heatmap of the number of differentially expressed genes (FDR < 0.05) detected between workers from the ten source colonies used to establish the microcolonies. We observed many differentially expressed genes between the 10 source colonies, with colonies two and three being particularly dissimilar. On average, we detected 445 differentially expressed genes between other colonies and colonies 2 or 3, compared to average 97 differentially expressed genes between other colony pairs. Importantly, microcolonies established using worker bees from each source colony were equally distributed among the treatments used. All colonies were purchased from a commercial breeder and were healthy. During the experiment all bees were kept under controlled conditions. Therefore, we expected the variation in gene expression to be driven by baseline-biological differences between the source colonies. We decided not to exclude colonies two and three from the analysis since the potential impact of these colonies on our results was mitigated by the presence of representative individuals from these source colonies in each treatment group.

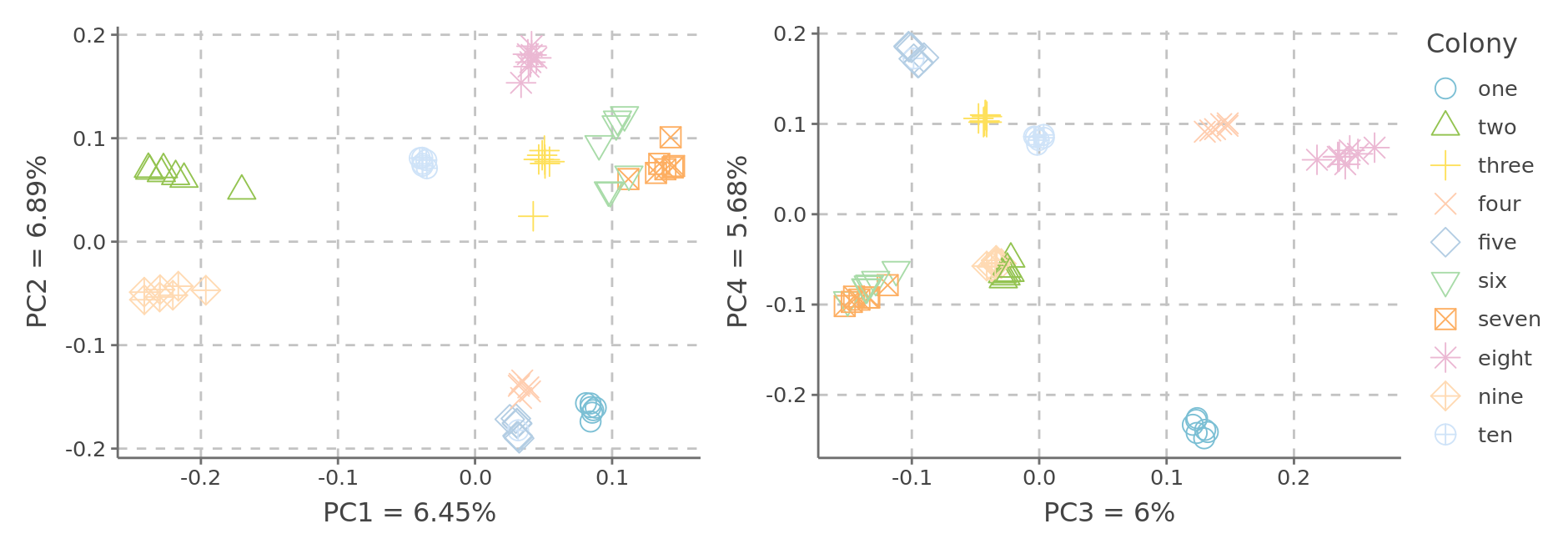

Figure S2. Principal component analysis of the SNPs detected between the 10 source colonies. Each data point indicates a pool of three workers from all microcolonies assembled using the corresponding source colony. All microcolonies used in the differential gene expression analysis were used here. All source colonies were clearly separated indicating underlying genomic differences that were controlled for by including the source colony in the DESeq2 model design.

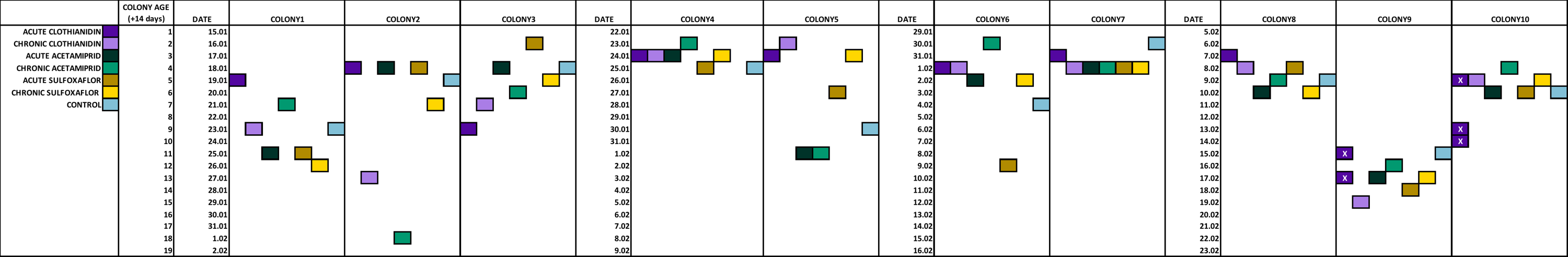

Figure S3. Experimental design. We obtained microcolonies from 10 source colonies and assigned to one of the seven treatments. All source colonies were two weeks old when we started arranging the microcolonies. We created microcolonies using callow workers. Workers within each microcolony emerged within 24h. Because of the differences in the pace of worker production, we created microcolonies in a staggered manner. The order in which we assigned treatments to the microcolonies was randomized. Our design ensured that treatments were assigned to microcolonies created at various stages of source colony development.
